# Local glucocorticoid signalling promotes pancreatic cancer

**DOI:** 10.1101/2025.11.14.688472

**Authors:** Clara Veiga-Villauriz, Hosni A.M. Hussein, Christopher J. Ward, Sanu K. Saji, Soura Chakraborty, Jhuma Pramanik, Yumi Yamashita-Kanemaru, Joanna Simpson, Sofia Laforest, Fede Diez, Natalie Z.M. Homer, Rahul Roychoudhuri, Timotheus Y. F. Halim, Bidesh Mahata

## Abstract

Pancreatic cancer (PC) represents one of the biggest challenges in terms of cancer treatment, mainly due to its continuously rising incidence, advanced stage at time of diagnosis, and dismal 5-year overall survival, which has not improved in recent decades despite the major advances made in oncological therapies. The limited progress in developing more effective therapies is, in part, attributable to the vast desmoplastic stroma present in PC. Additionally, immunosuppressive steroid-signalling has recently been shown to aid the development and metastasis of various tumour types. Therefore, we sought to explore whether local steroidogenesis and steroid signalling within the tumour microenvironment (TME) play a role in pancreatic cancer development. Reanalysis of publicly available datasets, including single cell RNA sequencing, as well as *in vivo* metastatic pancreatic ductal adenocarcinoma (PDAC) mouse models, allowed us to identify Hsd11b1 as the key enzyme responsible for locally elevated levels of the immunosuppressive glucocorticoid hormone, corticosterone. We identified fibroblasts as the major Hsd11b1-expressing populations in the pancreatic TME. Specifically, in mice, Hsd11b1 expression is primarily observed in iCAFs. Additionally, we found that patients with higher *HSD11B1* expression present an increased mortality rate as well as an enriched fibrotic signature and inhibited immune activity. Collectively, these findings suggest that *Hsd11b1* upregulation in iCAFs could be aiding PDAC development by promoting the activation of glucocorticoids directly in the TME. The presence of glucocorticoids inhibits inflammation and could also be enhancing local fibrosis by autocrine signalling in the fibroblast population. Given the urgent need for effective treatments in this fatal disease, targeting HSD11B1 represents a promising therapeutic strategy to overcome the immunosuppressive desmoplastic barrier and improve patient outcomes in pancreatic cancer.

## Introduction

Pancreatic cancer represents the third cancer-related cause of death globally with a 5% 5-year survival after diagnosis and is set to become the second leading cancer-related cause of death by 2030 in the US[1]. Most pancreatic cancer cases fall under the pancreatic ductal adenocarcinoma (PDAC) subtype, accounting for over 90% of diagnoses. This is partly due to the late stage at which most patients are diagnosed, due to the silent nature of the disease, but also because of its aggressiveness, chemoresistance, and general lack of response to immunotherapies^[2], [3]^. The limited progress in developing more effective therapies is, in part, attributable to the desmoplastic stroma, which hinders immune activation and restricts bloodstream access. The cellular stroma primarily consists of fibroblasts and immune cells, while the non-cellular part is largely a dense collagen matrix^[4], [5], [6]^.

Cancer associated fibroblasts (CAFs) are some of the key players involved in the exuberant production of extracellular matrix, collagen, and cytokines, leading to a fibroinflammatory stroma^[7], [8]^. Being some of the most abundant components within the PDAC TME, CAFs display a vast array of functions, both pro- and anti-tumorigenic, and are therefore considered a very heterogenous population. Advances in single cell technologies have allowed for their classification, primarily, into inflammatory CAFs (iCAFs), myofibroblastic CAFs (myCAFs), and antigen presenting CAFs (apCAFs)^[9], [10], [11], [12], [13]^.

In parallel, recent studies have shown that steroids can promote the development of several tumour types through immune-derived *de novo* steroidogenesis, particularly in those enriched with Th2 immune activity^[14], [15]^. Steroids are synthesised *de novo* from cholesterol and, through the process of steroidogenesis, can be further converted into a range of compounds that perform diverse physiological roles throughout the body. Cortisol and corticosterone are the most prevalent glucocorticoids (GCs) in humans and rodents, respectively, and play important roles in regulating metabolism, the stress response, or immune function. Their systemic release is controlled by the HPA axis and can be triggered by a range of different stressors^[16]^.

Alternatively, specific tissues, including tumour tissues, might increase their GC signalling by producing cortisol or corticosterone *in situ*^[17], [18], [19], [20], [21]^. Local GC production is attributed to the action of the enzyme 11β-hydroxysteroid dehydrogenase type 1 (HSD11B1). This enzyme catalyses the reduction of the 11-keto group in cortisone (in humans) or 11-dehydrocorticosterone (11-DHC, in rodents), generating the corresponding active 11β-hydroxylated glucocorticoid, cortisol or corticosterone, respectively. It is only capable of doing so when interacting with hexose-6-phosphate dehydrogenase (H6PDH), which provides the required NADPH co-substrate^[20], [22]^.

The role of HSD11B1-mediated steroid production in tumours, specifically PDAC, remains underexplored. Here, we investigated the expression of HSD11B1 in human PDAC tissues, as well as in primary and metastatic mouse tumours, and highlight iCAFs as the major producers. Additionally, we analyse the phenotypic differences between patients with varying levels of *HSD11B1* expression, finding that high expression of the enzyme correlates with enhanced fibrosis, more active fibroblasts, impaired immunity and exhausted CD8 T cells. For this reason, we identify *HSD11B1* as a potential bad prognostic factor in PDAC development and highlight it as a therapeutic target.

## Results

### HSD11B1 is locally upregulated in pancreatic cancer

To determine whether local activation of glucocorticoids (GCs) is taking place in pancreatic tumour tissue, we examined the expression of the enzyme HSD11B1 in human and mice. We accessed human transcriptomic and proteomic online databases through the GEPIA (Gene Expression Profiling Interactive Analysis) and UALCAN web platforms, which provide access to RNA-seq and proteomic datasets from TCGA and GTEx, and CPTAC, respectively. GEPIA analysis revealed that *HSD11B1* expression was significantly upregulated in PDAC tissues (TCGA, n = 179) when compared to normal tissue controls (GTEx, n = 171) as represented in Fig. 1A. In the same way, data from UALCAN (Fig. 1B) supported this finding at the proteomic level. Protein expression for HSD11B1 appears significantly higher in PDAC tissue samples (n = 157) when compared to normal tissue (n = 74). This was further supported by reanalysing Steele et al’s dataset (GSE155698) consistent of 16 PDAC and 3 adjacent/normal pancreatic tissue (from patients referred for Whipple procedures or distal pancreatectomy)^[23]^. Following quality control and normalisation, expression analysis of HSD11B1 demonstrated its restricted expression to the tumour site almost uniquely (Fig. 1C). Cells from the adjacent/normal pancreatic tissue exhibited reduced *HSD11B1* levels. Additionally, to mimic metastatic PDAC *in vivo,* we injected a Kras^LSL-G12D/+^;Trp53^LSL-R172H/+^;Pdx1-Cre;Rosa26^YFP/YFP^ murine pancreatic cancer cell line KPC-c3 intravenously (iv)^[24]^. Upon lung tumour development (KPC-c3 colonisation in the lung), immunofluorescence stains allowed us to corroborate that Hsd11b1 protein levels were indeed higher in tumour-bearing mice at 21d post-iv (Fig.1D). We corroborated this overexpression of Hsd11b1 in murine tumours by the integration and analysis of the following publicly available datasets: GSE165534^[25]^, GSE125588^[26]^, and GSE84133^[27]^. Indeed*, Hsd11b1* seems to be present at a higher level in the tumour samples when compared to normal pancreatic mouse tissue (Fig. 1E), supporting our human observation and our *in vivo* immunofluorescence data (Fig. 1D).

**Figure 1:**
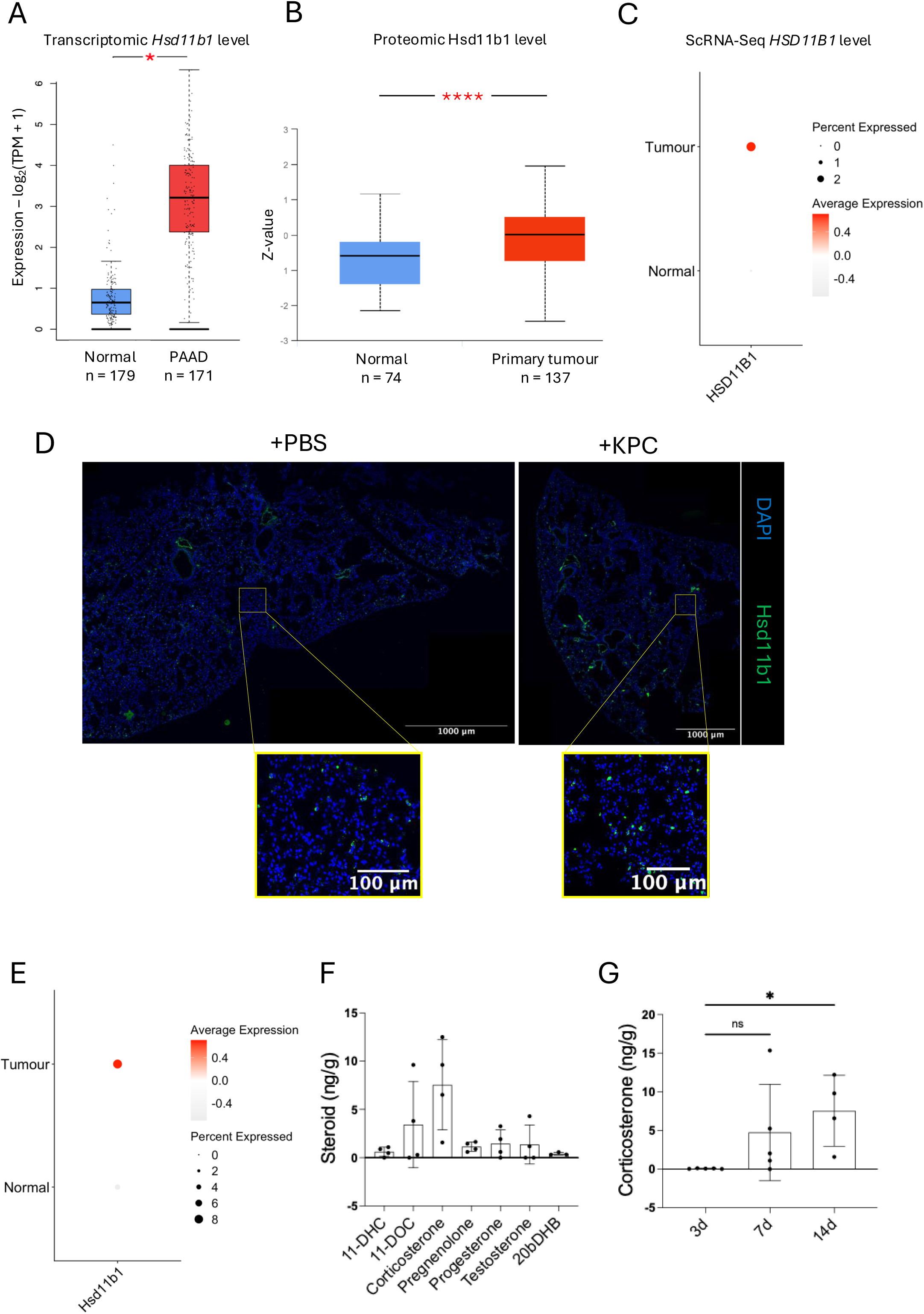
*Hsd11b1* is upregulated in human and murine pancreatic tumours, which results in increased local glucocorticoid presence. **A.** Boxplot representing *HSD11B1* expression in human pancreatic adenocarcinoma tumours (TCGA data; n=179) as compared to healthy tissue samples (GTEx data; n=171). Statistical comparisons were performed using standard unpaired Student’s t-test to assess diGerential gene expression. **B.** Proteomic data depicting HSD11B1 expression in human pancreatic adenocarcinoma (n=137) versus healthy samples (n=74) (CPTAC data). Statistical comparisons between groups are performed using Welch’s t-test to account for unequal variances and sample sizes. **C.** Reanalysed sc-RNASeq data showing the percentage of cells that express *HSD11B1* as well as its average expression in human pancreatic tumours (n=16) in comparison to adjacent normal pancreatic tissue (n=3). **D.** Representative immunofluorescence staining for Hsd11b1 in lung tissue from mice at 21d post-iv injection of KPC 2838c3 cells or PBS (vehicle). **E.** sc-RNASeq data depicting both the proportion of *Hsd11b1*-expressing cells and the average expression levels in mouse pancreatic tumours (n=13) compared to matched adjacent normal pancreatic tissue (n=2). **F.** LC-MS/MS quantification of steroids present in mouse lungs in a metastatic mouse model of pancreatic cancer (n=4). Steroid quantification was performed 14 days post intravenous injection of KPC 2838c3 cells. Bar plots ± SD. **G.** LC-MS/MS quantification of corticosterone at various timepoints post KPC 2838c3 intravenous injection (n = 4-5). Bar plots ± SD.

Furthermore, to attest for Hsd11b1 functionality, we aimed to quantify the levels of active corticosterone during tumorigenesis *in vivo* in mice. LC-MS/MS analysis of lung tissues from mice injected with the KPC-c3 pancreatic cancer cell line revealed high levels of corticosterone in the lungs of tumour-bearing mice (Fig.1F). In fact, corticosterone appears as the most highly expressed steroid (out of our panel) in the lungs at 14d post-iv injection (Fig. 1F). Furthermore, corticosterone levels in the lungs of tumour-injected mice increase steadily in parallel with tumour progression, and reached a significant level at 14d post-iv, underscoring the significant influence of Hsd11b1 activity on tumour development (Fig.1G).

### iCAFs are the key Hsd11b1-expressing cells in the PDAC TME

To further characterise the local steroid conversion in the PDAC TME we further investigated the expression of *HSD11B1* in the pancreatic stroma during tumour development. We performed unbiased clustering and dimensionality reduction through Uniform Manifold Approximation and Projection (UMAP), reanalysing Steele et al’s dataset^[23]^. 13 clusters of stromal PDAC cells were identified (Fig. 2A). The clusters were annotated based on lineage markers represented in Fig.2B. We identified the fibroblast population as the key *HSD11B1*-expressing population in the human tumour (Fig. 2C). When comparing *HSD11B1* expression in CAFs with fibroblasts from adjacent normal tissue, we observe that tumour-associated fibroblasts show both a higher expression level (Fig.2D) as well as a significantly greater proportion of *HSD11B1*-positive cells among all fibroblasts (Fig.2E).

**Figure 2:**
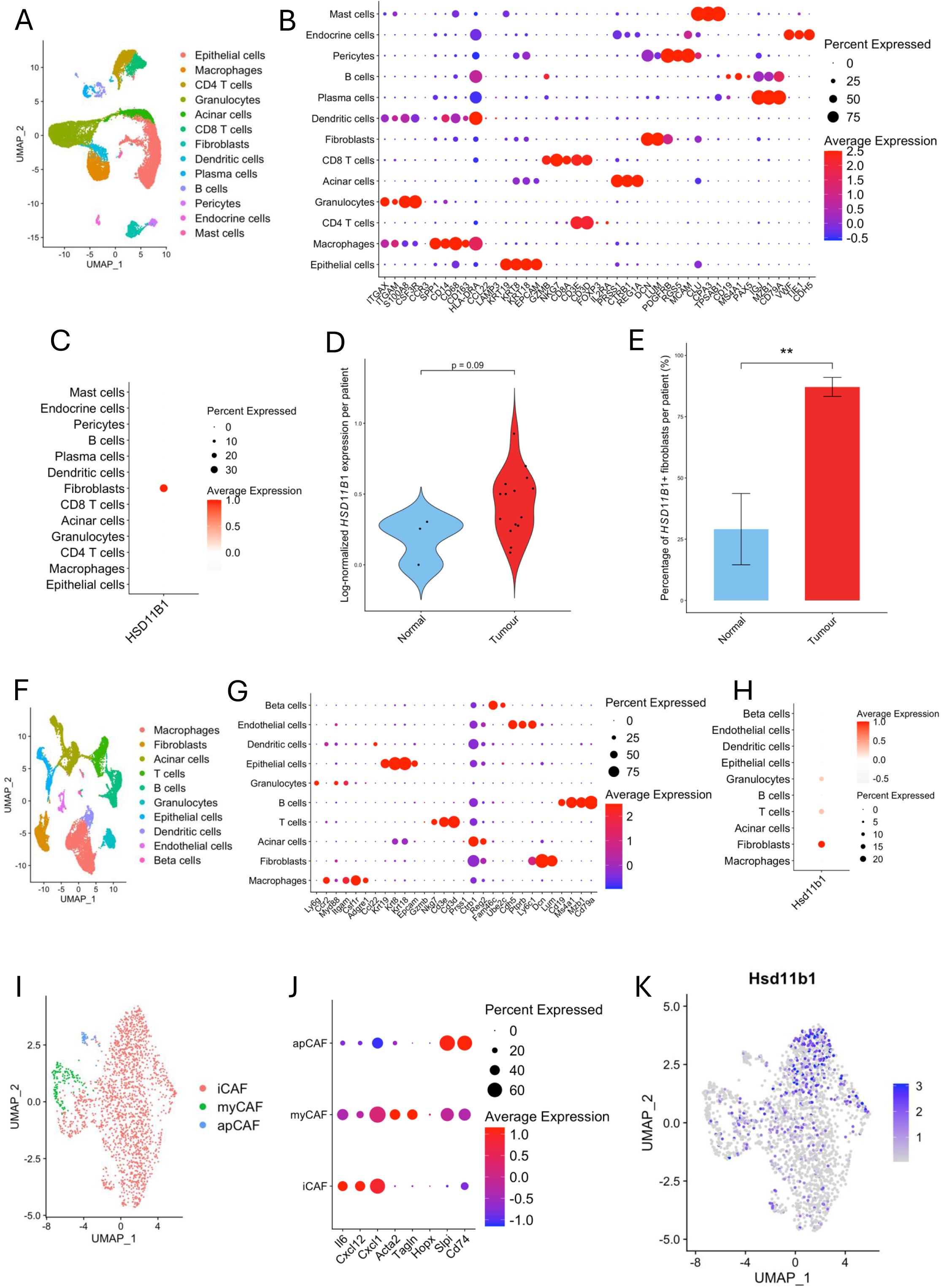
*Hsd11b1* upregulation is limited to fibroblasts, particularly to the iCAF subtype, in pancreatic ductal adenocarcinoma. A. UMAP on 13 cell populations distinguished in 16 human PDAC tumour samples obtained from publicly available data (GSE155698). B. Dot plot of the markers used to identify the cell populations contained in the tumour samples. The percentage of cells expressing each marker as well as the average expression are depicted in the form of a colour gradient and dot size, respectively. C. Single-cell transcriptomics data showing *HSD11B1* expression in each of the cell populations identified in human PDAC tissue. D. *HSD11B1* expression per patient in fibroblasts from adjacent normal tissue (n=3) versus CAFs in tumour samples (n=16). E. Percentage of fibroblasts expressing *HSD11B1* per patient in normal tissue (n = 3) versus tumour tissue (n = 16). F. Single-cell transcriptomics data showing *Hsd11b1* expression in each of the cell populations identified in mouse PDAC tissue. G. Markers used to identify cell populations in tumour and normal mouse samples. Dot size represents the percentage of cells expressing each marker and colour indicates average expression. H. *Hsd11b1* expression in the diGerent identified mouse tumour tissue cell clusters. I. UMAP representation of the three identified CAF populations within the fibroblast cluster from tumour samples. J. Markers used to identify each of the CAF populations, iCAF, myCAFs and apCAFs. The percentage of cells expressing each marker as well as the average expression are depicted in the form of a colour gradient and dot size, respectively. K. Single-cell transcriptomic analysis depicting *Hsd11b1* expression across the distinct murine CAF subpopulations.

In order to corroborate these findings in murine models, we reanalysed the GSE165534, GSE125588, and GSE84133 datasets^[25], [26], [27]^. Upon integration, we performed unbiased clustering and dimensionality reduction through UMAP, leading to the identification of 10 cell clusters (Fig. 2F). The clusters were annotated based on murine lineage markers represented in Fig. 2G. We found that in murine tumours *Hsd11b1* is expressed predominantly in fibroblasts. We also observed T cell and granulocyte expression of the enzyme, albeit lower lever compared to fibroblasts (Fig. 2H).

We further reclustered this murine dataset to examine *Hsd11b1* expression in the three major murine CAF subtypes, iCAFs, myCAFs, and apCAFs (Fig.2I). Each cluster was annotated based on lineage markers represented in Fig.2J. We identified iCAFs as the major *Hsd11b1-*expressing CAF subtype in murine PDAC (Fig. 2K).

### HSD11B1 expression impacts patient survival rate, probably due to impaired immune function and exacerbated fibrosis

The elevated HSD11B1 expression observed in the PDAC microenvironment likely translates to higher cortisol concentrations in human tumours. Therefore, we hypothesised that patients with higher levels of HSD11B1 would show signs consistent with prolonged steroid signalling that could affect their survival or disease progression. To determine the effect of *HSD11B1* expression in patient outcome, we accessed TCGA survival data. Additionally, to better characterise the differences between patients expressing high versus low levels of *HSD11B1*, we reanalysed the previously described dataset compiled by Steele et al, 2021^[23]^.

To assess the association between *HSD11B1* expression and overall survival in PDAC, patients were stratified using two different expression cutoffs. First, we divided the cohort at the median *HSD11B1* expression level. Although the Kaplan–Meier curves showed a visible separation between high and low expressors (Fig. 3A), the difference did not reach statistical significance, potentially indicating that a median-based division may not fully capture the prognostic impact of *HSD11B1* expression. We then applied a more selective stratification by isolating the lowest 25% of *HSD11B1* expressors as the “low” group and comparing them to the remaining 75% of patients. This analysis produced a more pronounced divergence between the survival curves, with the low-expression group exhibiting noticeably reduced overall survival (Fig. 3B). These findings indicate that the survival difference becomes more apparent when focusing on patients with the most markedly reduced *HSD11B1* expression levels.

**Figure 3:**
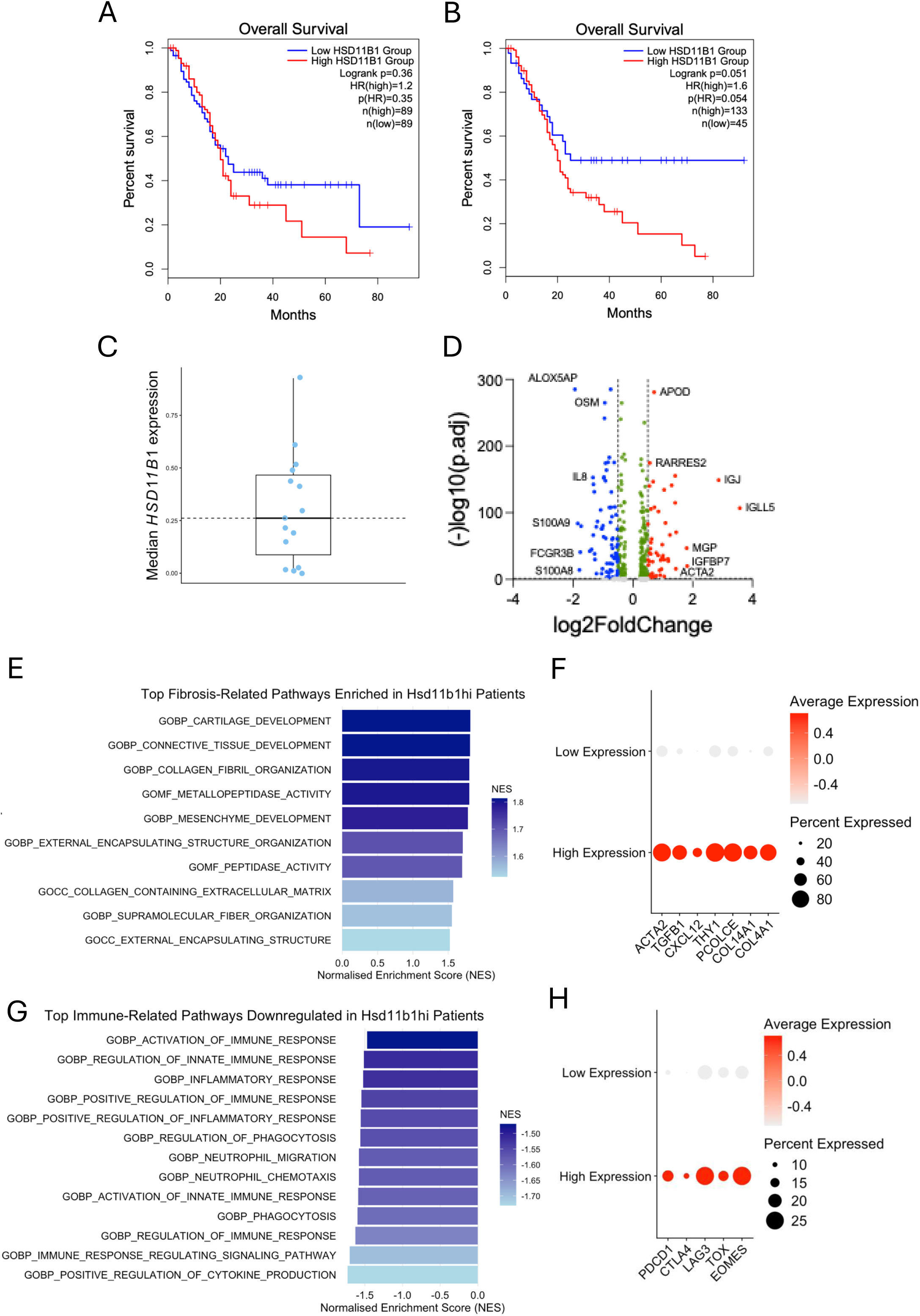
High levels of *HSD11B1* expression in PDAC patients correlates with reduced survival, defective immunity, and enhanced fibrosis. **A.** Kaplan-Meier survival curve depicting the survival probability of patients stratified by *HSD11B1* expression levels, with groups defined as above or below the median expression value. The data is sourced from the TCGA dataset using the GEPIA webpage. **B.** Kaplan-Meier survival curve depicting the survival probability of patients stratified by *HSD11B1* expression levels, with groups defined as the lowest (<25%) and highest (<25%) expression values. The data is sourced from the TCGA dataset using the GEPIA webpage. **C.** Box plot showing the variety of median *HSD11B1* expression levels within the various patient fibroblast’s tumour samples from the GSE155698 publicly available dataset. **D.** Volcano plot showing diGerentially expressed genes in CAFs from *HSD11B1^hi^* and *HSD11B1^low^* patients. Each dot represents a gene. **E.** GSEA analysis showing the hallmark biological processes related to fibrosis enriched in *HSD11B1^hi^* patients with normalised enrichment scores (NES) represented in the x axis. **F.** Single-cell transcriptomics data showing the expression of fibrotic marker genes in isolated CAFs from high *HSD11B1*-expressing patients versus low. **G.** GSEA analysis depicting the top immune-related biological processes downregulated in HSD11B1^hi^ patients with normalised enrichment scores (NES) represented in the x axis. **H.** Single-cell transcriptomics data showing the expression of T cell exhaustion genes in isolated tumour infiltrating CD8 T cells from high *HSD11B1*-expressing patients versus low.

To further explore this, we categorised the 16 patients from the Steele et al, 2021 study in the same manner, above or below the median *HSD11B1* expression (Fig. 3C). Differential gene expression analysis (DGE) identified 298 significantly regulated genes (151 upregulated in red, and 147 downregulated in blue) (Fig. 3D). Notably, among the strongly induced genes were *RARRES2, ACTA2, MGP,* and *IGFBP7*, all previously linked to fibrotic-like processes (Fig. 3D; red). Alternatively, genes like *IL8, OSM, ALOX5AP,* and *S100A8/9* were downregulated, all involved in the regulation of inflammatory and immune responses (Fig.3D; blue). Gene set enrichment analysis (GSEA) identified 76 pathways differently regulated in both patient cohorts. A high number of the enriched pathways were associated with fibrosis-related processes, which are represented in Fig. 3E. Indeed, fibroblasts isolated from *HSD11B1*^hi^ patients present elevated levels of some fibroblast activator/fibrotic markers in their fibroblasts (Fig. 3F). Meanwhile, most downregulated pathways involved processes of immune activation and other inflammatory responses (Fig. 3G). Interestingly, CD8 T cells present in patients with high *HSD11B1* expression express higher levels of T cell exhaustion markers, known to be elevated upon steroid exposure and signalling (Fig.3H). These data suggest the strong correlation of *HSD11B1* presence with a fibrotic signature and dysregulated immunity.

## Discussion

This study demonstrates that HSD11B1 is significantly upregulated in pancreatic ductal adenocarcinoma (PDAC) and establishes inflammatory cancer-associated fibroblasts (CAFs) as the primary cellular source of local glucocorticoid production within the tumour microenvironment. Elevated HSD11B1 expression correlates with a distinct pathological phenotype characterized by enhanced fibrosis and impaired immune function, providing mechanistic insight into how local steroid metabolism facilitates immune evasion in PDAC. The immunosuppressive properties of glucocorticoids are well established, and recent work has highlighted the importance of tumour-intrinsic steroidogenesis in promoting immune escape across multiple malignancies^[21], [28], [29], [30]^. While previous studies have focused primarily on *de novo* steroidogenesis mediated by CYP11A1 in T cells and macrophages^[14], [15], [31]^, our data identify HSD11B1-mediated metabolite recycling as an alternative mechanism through which the PDAC microenvironment amplifies local glucocorticoid signalling. Unlike the synthesis of steroids from cholesterol, HSD11B1 rapidly converts inactive cortisone or 11-dehydrocorticosterone into active cortisol or corticosterone, in human or rodents respectively^[20]^, providing efficient elevation of glucocorticoid levels at the tumour site. Mass spectrometry analysis confirmed significantly elevated corticosterone levels in murine metastatic tumours, establishing functional relevance for HSD11B1 activity in cancer progression.

Single-cell transcriptomics analyses revealed that fibroblasts represent the predominant *HSD11B1*-expressing population in human PDAC, with enrichment specifically within the inflammatory CAF (iCAF) subset in murine models. This finding is mechanistically coherent given that iCAFs emerge in response to inflammatory signals such as IL-1a^[9]^. IL1a, among other inflammatory cytokines, is also known to induce *HSD11B1* expression in other contexts^[19], [32], [33]^. We propose that inflammation-driven iCAF activation triggers local glucocorticoid production, via Hsd11b1 upregulation, to dampen immune responses. Although we detected *Hsd11b1* expression in T cells, macrophages, and granulocytes within murine tumours, fibroblasts remained the dominant source in human pancreatic cancer. Interestingly, cancer cells did not emerge as significant *HSD11B1* expressors in our analyses, contrasting with recent reports demonstrating functional HSD11B1 expression in tumour cells from other cancer types^[21], [34]^. This discrepancy may reflect tumour-specific differences in cellular steroidogenic capacity.

Patients with high HSD11B1 expression exhibited a transcriptomic signature characterised by elevated fibrosis-related pathways and suppressed immune activation. Fibroblasts from these patients expressed increased levels of fibrotic markers including *Col14a1, Col4a1, Pcolce, Thy1, Cxcl12, Acta2, Tgfb1, and Il1b,* while CD8 T cells displayed an exhaustion phenotype consistent with prolonged glucocorticoid exposure. These observations are particularly relevant given that the desmoplastic stroma represents a major barrier to both immune infiltration and therapeutic efficacy in PDAC. While the relationship between glucocorticoids and fibrosis remains complex and context-dependent across different tissues and disease states^[35], [36], [37], [38], [39], [40], [41], [42]^, our data support a model in which HSD11B1-mediated glucocorticoid production contributes to both the immunosuppressive and desmoplastic features of pancreatic cancer. Recent work from our laboratory has elucidated the molecular mechanisms through which glucocorticoids cooperate with RUNX transcription factors to drive CD8 T cell dysfunction^[43]^, providing a direct mechanistic link between HSD11B1-derived glucocorticoids and the exhausted T cell phenotype observed in HSD11B1-high patients. Furthermore, genetic ablation of HSD11B1 in cancer cells has been shown to enhance immune activity in preclinical models, supporting the therapeutic rationale for targeting this enzyme^[21], [34]^.

To evaluate the prognostic relevance of *HSD11B1* expression in pancreatic cancer, we examined patient survival using two different stratification approaches. When the cohort was divided at the median expression level, the resulting Kaplan–Meier curves showed a visible separation between high and low expressors, although the difference did not reach statistical significance. This suggests that *HSD11B1* expression may influence patient outcome; however, a median cutoff may not adequately capture biologically meaningful variation in enzyme activity.

We hypothesize that the relationship between enzyme expression and glucocorticoid-generating capacity may involve a threshold effect, such that only markedly reduced expression meaningfully impairs enzymatic activity and thus cortisol production. If this is the case, patients with expression levels near the median may retain sufficient enzyme function to produce similar glucocorticoid signalling and downstream biological effects. Consequently, dividing patients at the median could obscure survival differences that are most apparent among those with substantially lower expression.

Future studies should address several key questions to establish the therapeutic potential of HSD11B1 inhibition. First, direct steroid profiling of primary human pancreatic tumours is necessary to quantify glucocorticoid levels *in situ* and confirm functional enzyme activity in the clinical context. Second, functional validation through genetic or pharmacological HSD11B1 inhibition in primary PDAC models is required to establish causality and assess therapeutic efficacy. Finally, the mechanistic interplay between HSD11B1-derived glucocorticoids and fibroblast activation warrants investigation, particularly potential autocrine effects on CAF differentiation and extracellular matrix production. Selective targeting of the immunosuppressive functions of iCAFs through HSD11B1 inhibition may offer a refined therapeutic strategy that preserves beneficial stromal functions while restoring anti-tumour immunity.

In conclusion, we identify HSD11B1 as a key regulator of local glucocorticoid production in pancreatic ductal adenocarcinoma, with iCAFs serving as the principal cellular source. High HSD11B1 expression correlates with a fibrotic, immunosuppressive tumour microenvironment and warrants further investigation as a therapeutic target to overcome immune evasion in this defiant malignancy.

## Materials and methods

### Patients

This study used publicly available *HSD11B1* gene expression data from 179 patients with PAAD (Pancreatic Adenocarcinoma) from the TCGA dataset and 171 normal pancreatic tissue samples from GTEx. Both datasets were accessed through the online tool GEPIA2, which normalises the expression levels across TCGA and GTEx samples, a pipeline based on that of the Xena project (https://xenabrowser.net/)^[19], [44]^. Briefly, Raw RNA-seq data from both sources are normalised using the TPM method and following log2 transformation to stabilise variance and resolve the differences in sequencing depth. Protein levels for HSD11B1 were obtained from UALCAN, where the proteomic data is derived from mass spectrometry global proteome profiling of primary tumour matched to normal tissue^[45]^ (https://ualcan.path.uab.edu). A total of 137 PAAD tumours and 74 paired adjacent non-tumour controls were accessed.

### Single-cell RNA sequencing, unsupervised clustering, and visualisation

The previously published and publicly available single cell RNA sequencing (scRNA-Seq) datasets used in this study were as follows: GSE165534^[25]^, GSE125588^[26]^, GSE84133[27] and GSE155698^[23]^. The data was downloaded from NCBI GEO (https://www.ncbi.nlm.nih.gov/geo/) and processed with the standard scRNA-seq integration workflow in Seurat (v4.4.0) to mitigate batch effects. Cells displaying a gene count greater than 200 but fewer than 2,500 as well as less than 5% of their genes as mitochondrial were retained.

For clustering, the 2,000 most variable genes were selected using the *FindVariableFeatures* function in Seurat. Principal component analysis (PCA) was subsequently performed, and dimensionality reduction for visualization was achieved using Uniform Manifold Approximation and Projection (UMAP). The selection of significant principal components (PCs) for downstream analysis was guided by Seurat’s *ElbowPlot* function, resulting in the identification of 10 PCs. Cell clustering was conducted in PCA space using the Shared Nearest Neighbor (SNN) algorithm, implemented via the *FindNeighbors* and *FindClusters* functions in Seurat v3. The resulting clusters were visualized in UMAP space using the DimPlot function and annotated by finding the differentially expressed genes in each cluster using *FindAllMarkers* in Seurat. Clusters were then identified by previous annotation and confirmed using Cluster Identity Predictor (CIPR). Gene expression patterns were visualized using the ggplot2 package (v4.0.0).

### Pathway analysis

Patients were stratified into two groups, HSD11B1^hi^ and HSD11B1^low^, based on the median expression level of *HSD11B1*. Differentially expressed genes (DGE) between groups were identified using the *FindMarkers* function. The volcano plot was generated in GraphPad Prism (v10.4.2) to visualise gene expression differences. Gene Set Enrichment Analysis (GSEA) was performed using the fgsea package in R (v1.34.2). Genes were ranked according to the metric *sign*(log₂ fold change) × –log₁₀(*P* value), thereby incorporating both effect size and statistical significance. Enrichment was tested against the human hallmark gene sets C5 (Ontology gene sets) from the Molecular Signatures Database (MsigDB), followed by filtering of results for p values <0.05, and visualisation in graphical formats. All steps were performed in R (4.5.1).

### Mice

Animal experiments were conducted in accordance with UK Home Office guidelines, under the Project License P0AB4361E or PP2448972, and approved by Cambridge University Biomedical Services Gurdon Institute’s Animal Welfare Ethical Review Body. All mice were housed following the UK Animals (Scientific Procedures) Act 1986 and its Amendment Regulations 2012 guidelines. The animal facility was maintained under a 12h light/dark cycle, in a controlled ambient temperature of 21 °C ± 2 °C, and relative humidity between 45% and 65%. Animals were housed in individually ventilated cages, grouped to a maximum of five animals per cage. Environmental enrichment such as tunnel, gnawing, and nesting material was present, and a standard diet was provided. Mice were sex- and age-matched whenever possible, and most mice were used at 8 to 12 weeks of age.

### Tumour metastasis model

Mouse pancreatic Kras^LSL-G12D/+^;Trp53^LSL-R172H/+^;Pdx1-Cre;Rosa26^YFP/YFP^ (KPCY)-derived 2838c3 cancer cell line, KPC-c3^[24]^ were passaged in DMEM (Sigma, D6429) supplemented with 5% Heat-Inactivated FBS (Sigma, F9665) and antibiotics (Sigma, P4333). To generate the metastatic lung tumours, mice were injected intravenously with 0.5x10^6^ KPC-c3 cells in sterile 100ul PBS. Control animals were injected with 100ul sterile PBS. Lungs were dissected at 3-, 7-, 14-, or 21-days post-injection.

### Confocal Immunofluorescence Staining

Frozen tissue sections (7um) were fixed in 10% formalin solution (Sigma, HT5011) and permeabilised using 0.2% triton-x. Staining was performed with a primary antibody against Hsd11b1 (1:100, ThermoFisher, PA5-21586) overnight at 4 °C. Sections were then stained with a goat-anti Rabbit CoraLite Plus 555 secondary antibody (1:200, Proteintech, RGAR003). Images were acquired using a Leica SP8 confocal at 20x magnification and analysed using ImageJ2 (v2.14). All images were background-subtracted ahead of further analysis.

### Steroid Quantification via Liquid Chromatography Mass Spectrometry (LC-MS/MS)

Lung tissue (∼ 25mg) from either tumour-challenged mice or PBS controls was placed in 2 mL reinforced tubes containing 1.4 mm ceramic beads (FisherScientific). 1 mL of acetonitrile with 0.1% formic acid (v/v) and 20 µL of an isotopically labelled steroid standard mixture were added to the tube. Samples were homogenized at 1 m/s for 30 seconds for 3 cycles using a Bead Ruptor 24 Elite (Omni International) equipped with a CryoCool unit. Supernatants were transferred to a Filter+ plate (Biotage, Sweden), and eluate collected into clean 96-well plates. The filtered homogenate underwent further processing through phospholipid depletion (PLD+) plates (Biotage, Sweden). Eluate was dried, reconstituted in water/methanol (70:30 v/v), and sealed with zone-free plate seals, ready for LC-MS/MS analysis.

An I-Class UPLC (Waters, UK) with a Kinetex C18 column (150 × 2.1 mm, 2.6 μm) was used for liquid chromatography with a mobile phase of water and methanol, both containing 0.05 mM ammonium fluoride, starting at 50% methanol (B), increasing to 95%, then returning to 50%. The flow rate was 0.3 mL/min. The column was maintained at 50°C and the autosampler at 10°C, with a 20 µL injection volume. Each analytical run lasted 16 minutes per sample. Steroid detection was performed using a QTRAP 6500+ mass spectrometer (AB Sciex, Warrington, UK) equipped with an electrospray ionization Turbo V ion source, using method described previously^[46]^. The ion spray voltage was set at +5500 V for positive mode and -4500 V for negative mode, with a source temperature of 600°C. Multiple reaction monitoring parameters were optimized for each steroid, including pregnenolone (P5) with transitions at *m/z* 317.1→281.1 and *m/z* 317.1→159.0, and its labelled standard (^13^C_2_,d2-P5) at *m/z* 321.2→285.2. Parameters such as declustering potential (DP), collision energy (CE), and collision cell exit potential (CXP) were set accordingly, with retention times around 10.4 minutes. P5 concentrations were quantified by calculating the ratio of analyte (P5) to its labelled standard (^13^C_2_,d2-P5) peak areas, followed by linear regression analysis using MultiQuant 3.0.3 (AB Sciex, UK). This approach was applied to other steroids including corticosterone, 11-dehydrocorticosterone, progesterone, and testosterone.

### Statistics

Statistical analyses were performed using Graphpad Prism (v10.4.2). Two-tailed Student’s t-tests or Kruskal-Wallis test with Dunn’s for multiple comparisons were applied as appropriate, with p-values < 0.05 considered statistically significant. For online available data accessed via GEPIA2 or UALCAN, the platforms’ built-in statistical tests were applied and are specified in the figure legends. For DEG analysis in single-cell datasets, p values were computed using the default methods implemented in the corresponding R packages.

For all experiments, N defines the cohort size or biological replicates. Each representative experimental result displayed represents at least two independent experiments. Steroid profiling (LC-MS/MS) experiments were performed once with independent biological replicates.

## Resource availability Lead contact

Further information and requests for resources and reagents should be directed to and will be fulfilled by the Lead Contact, Bidesh Mahata

## Material availability

This study did not generate unique reagents.

## Data and code availability

Data: No new large datasets (e.g., RNA-seq) generated in this study. Code: This paper does not report original code.

Additional information: Any additional information required to reanalyse the data reported in this paper is available from the lead contact upon request.

## Author contribution

CVV: Acted as project lead, conceptualised, designed and performed the experiments, analysed, interpreted & visualised the data. Wrote the manuscript. HAMH: Aided experimental design and study conceptualisation. Contributed to mice experiments, tissue collection and processing. CJW: Contributed to *in vitro* experiment design and work. SKS: Contributed to mice experiments, tissue collection, and processing. SC: Supported scRNA-Seq analysis. JP, YYK: Contributed to mouse experiments. FD, JS and SL: Extracted and analysed samples by LC-MS/MS. NZMH: Designed the experiments for steroid analysis and interpreted mass spectrometry data. KO: Supervision and *in vivo* study aid. TYFH: conceptualised and supervised the study. BM: conceptualised and supervised the study. All authors read and approved the draft manuscript before submission.

## Acknowledgements

We appreciate the support and animal husbandry provided by the UBS animal facility at the Gurdon Institute. We also thank Prof Klaus Okkenhaug for his mentorship throughout the project and support with *in vivo* experiments. We are grateful to Scott Denham in the Edinburgh Clinical Research Facility, University of Edinburgh for his technical expertise in operation of the LC-MS/MS. We would also like to thank the Cambridge Advanced Imaging Centre (CAIC) for their assistance. We used BioRender.com to generate the graphical illustrations presented in this manuscript.

## Data and materials availability

All newly generated data are presented/displayed in the article.

